# SARS-CoV-2 Variants Reveal Features Critical for Replication in Primary Human Cells

**DOI:** 10.1101/2020.10.22.350207

**Authors:** Marie O. Pohl, Idoia Busnadiego, Verena Kufner, Irina Glas, Umut Karakus, Stefan Schmutz, Maryam Zaheri, Irene Abela, Alexandra Trkola, Michael Huber, Silke Stertz, Benjamin G. Hale

## Abstract

Since entering the human population, SARS-CoV-2 (the causative agent of COVID-19) has spread worldwide, causing >100 million infections and >2 million deaths. While large-scale sequencing efforts have identified numerous genetic variants in SARS-CoV-2 during its circulation, it remains largely unclear whether many of these changes impact adaptation, replication or transmission of the virus. Here, we characterized 14 different low-passage replication-competent human SARS-CoV-2 isolates representing all major European clades observed during the first pandemic wave in early 2020. By integrating viral sequencing data from patient material, virus stocks, and passaging experiments, together with kinetic virus replication data from non-human Vero-CCL81 cells and primary differentiated human bronchial epithelial cells (BEpCs), we observed several SARS-CoV-2 features that associate with distinct phenotypes. Notably, naturally-occurring variants in Orf3a (Q57H) and nsp2 (T85I) were associated with poor replication in Vero-CCL81 cells but not in BEpCs, while SARS-CoV-2 isolates expressing the Spike D614G variant generally exhibited enhanced replication abilities in BEpCs. Strikingly, low-passage Vero-derived stock preparation of 3 SARS-CoV-2 isolates selected for substitutions at positions 5/6 of E, and were highly attenuated in BEpCs, revealing a key cell-specific function to this region. Rare isolate-specific deletions were also observed in the Spike furin-cleavage site during Vero-CCL81 passage, but these were rapidly selected against in BEpCs, underscoring the importance of this site for SARS-CoV-2 replication in primary human cells. Overall, our study uncovers sequence features in SARS-CoV-2 variants that determine cell-specific virus replication, and highlights the need to monitor SARS-CoV-2 stocks carefully when phenotyping newly emerging variants or potential variants-of-concern.

## INTRODUCTION

Severe Acute Respiratory Syndrome Coronavirus 2 (SARS-CoV-2), a novel betacoronavirus belonging to the *Coronaviridae* family, appears to have first entered humans in late 2019 in the Hubei province of China [1]. Since then it has spread worldwide in the human population, predominantly causing a mild-to-severe respiratory disease, termed COVID-19. Currently, pandemic SARS-CoV-2 has led to more than 100 million laboratory-confirmed COVID-19 cases globally, resulting in more than 2 million deaths [2].

Coronaviruses are enveloped viruses with large positive-sense RNA genomes of ∼30kb [3]. The SARS-CoV-2 Spike (S) glycoprotein resides on the surface of virions, and mediates viral entry into the host cell by binding to cellular ACE2 [4, 5] and triggering viral-host membrane fusion [3, 6]. This fusion function of S is dependent on its cleavage by host cell proteases, which occurs either following attachment of virions to the host cell membrane or during virion maturation and egress [3, 7, 8]. Strikingly, SARS-CoV-2 S harbors a furin-cleavage site [9], and pre-activation ‘priming’ of S by furin-like proteases increases fusion-dependent entry efficiency of the virus at the plasma membrane following further cleavage by host TMPRSS2 [10-13]. Other SARS-CoV-2 virion components include the Membrane (M) protein that confers shape and support to the virus particle and interacts with both S and the Nucleoprotein (N; which coats the viral RNA genome [14, 15]), and the Envelope (E) protein, a small membrane protein that both drives virion assembly and budding [16], and has ion channel activity linked to viral pathogenesis [17].

Following S-mediated entry, translation of the viral genomic RNA results in expression of several non-structural proteins (nsp1-16), many of which form the essential replicase-transcriptase complex or fulfill additional important functions in the virus life cycle, such as evasion of host innate immunity [3]. Notably, the replication complex of coronaviruses includes proofreading activity, leading to greater genome stability as compared to other RNA viruses that typically lack this function [18]. Nevertheless, several mutations in the SARS-CoV-2 genome have already been reported in human-circulating viruses worldwide, and some exhibit increasing prevalence suggestive of positive-selection or fitness advantage in the new human host [19-21]. An important example of this is the D614G substitution in S, which rapidly dominated SARS-CoV-2 sequences following its emergence, and is functionally linked to increased virus infectivity and replication capacity [22-25], potentially aiding virus spread throughout the population. Other recently acquired mutations in the SARS-CoV-2 genome also appear to confer an advantageous replication and/or transmission phenotype in humans, and may impact virus antigenicity and the effectiveness of deployed vaccines [21].

In this study, we characterized 14 SARS-CoV-2 isolates from patient material collected during the first pandemic wave in Switzerland between March and May 2020. Phylogenetically, the isolates were representative of the different virus clades circulating in Europe in early 2020 [26]. While the individual isolates each carried distinct genetic variants that were already detected in the original patient material, rare additional mutations (notably in S and E) were observed upon low-passage virus stock preparation in Vero-CCL81 cells. Comparative replication and passage analysis of each SARS-CoV-2 isolate in Vero-CCL81 cells and differentiated primary human bronchial epithelial cells revealed the critical importance of the S furin-cleavage site and E positions 5/6 as specific determinants of SARS-CoV-2 infectivity in human respiratory cells. Furthermore, we observed that SARS-CoV-2 isolates expressing S G614 generally appeared to replicate more efficiently than isolates harboring S D614. Notably, SARS-CoV-2 isolates with naturally-occurring Orf3a (Q57H) and nsp2 (T85I) substitutions exhibited a cell-specific replication phenotype, with efficient propagation restricted to primary human bronchial epithelial cells. Overall, our comparative functional and sequence characterization of a broad range of virus isolates describes both important conserved residues, as well as naturally-occurring substitutions, that contribute to efficient SARS-CoV-2 replication in human respiratory cells.

## RESULTS

### Isolation and Sequence Analysis of SARS-CoV-2 from Patient Diagnostic Samples in Switzerland, March-May 2020

Following identification of the first SARS-CoV-2 positive patient in Switzerland at the end of February 2020, the initial pandemic wave was characterized by a peak of >1,000 new confirmed cases per day by mid-March, and a slow decline to a relatively stable ∼50 new cases per day by mid-to-late May (**Figure 1A**). From a biobank of over 1650 SARS-CoV-2 RT-PCR positive patient samples collected for diagnostic purposes throughout this period, we attempted virus isolations from 67 independent samples. Isolation success could readily be stratified by RT-PCR cycle threshold (Ct) value: we were able to isolate 62% of SARS-CoV-2 isolates with initial diagnostic Ct values <25 (n=21), while SARS-CoV-2 was only isolated from a single sample with diagnostic Ct values >25 (n=46) (**Figure 1B and Supplementary Table S1**). These isolation data are broadly in-line with SARS-CoV-2 culture success rates and correlations with Ct values described by others [24, 27-29], and indicate that presence of cell-culture infectious virus can be predicted from initial diagnostic RT-PCR results.

**Fig. 1.**
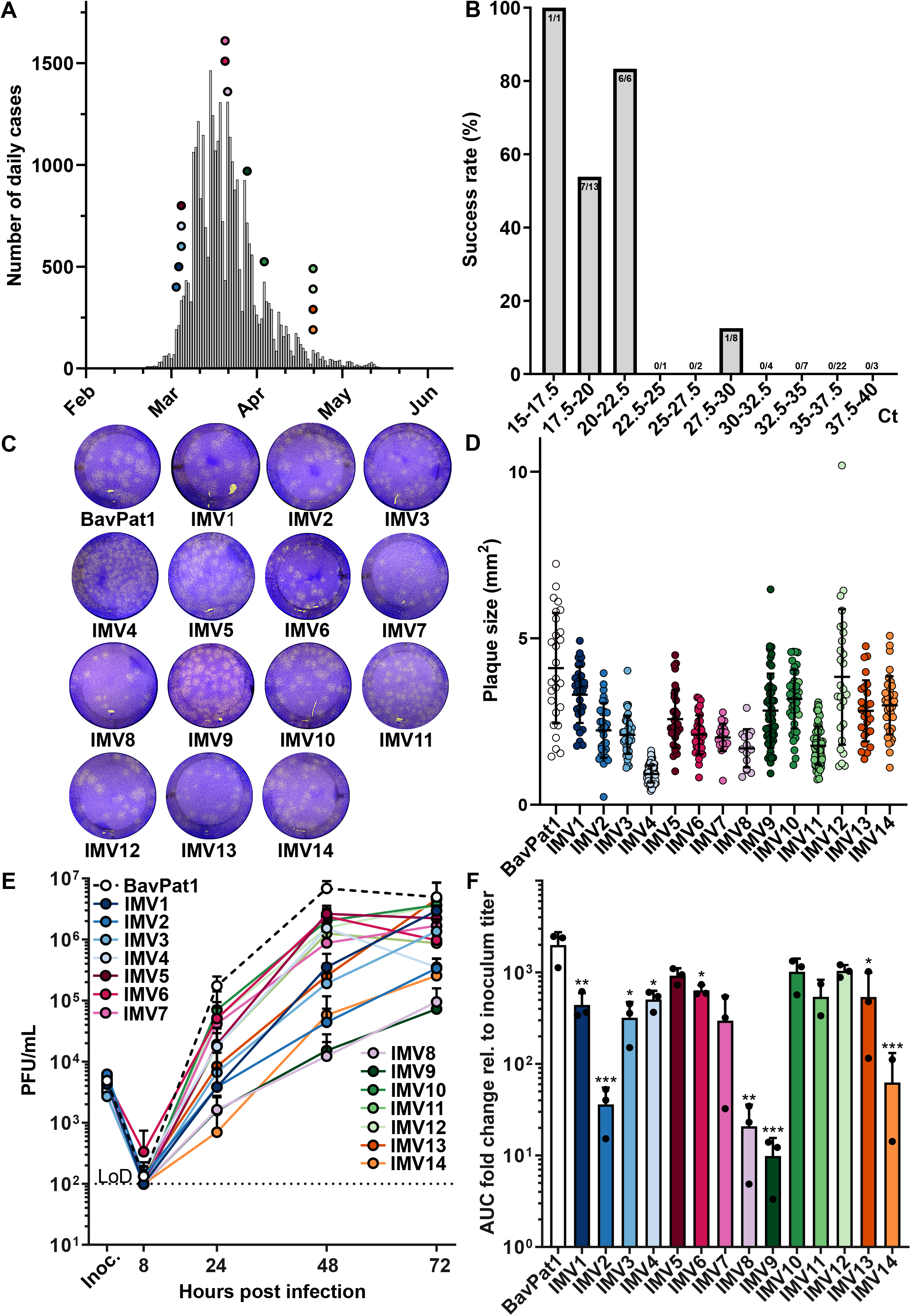
Isolation of SARS-CoV-2 from Patient Diagnostic Samples and Characterization in Vero-CCL81 Cells. (**A**) SARS-CoV-2-positive case counts per day in Switzerland between 24^th^ February 2020 and 31^st^ May 2020 (Source: Swiss Federal Office of Public Health - www.bag.admin.ch). Circles depict the day of patient sample acquisition of the 14 SARS-CoV-2 isolates generated in this study. (**B**) SARS-CoV-2 isolation success rate stratified by initial diagnostic Ct value of patient material. 67 PCR-positive patient samples were subjected to virus isolation attempts on Vero-CCL81 cells. Positive virus cultures exhibited cytopathic effect and low SARS-CoV-2-specific Ct values (<22) after 4-6 days of culture. Numbers at the top of bars represent isolation success/attempts. Numbers on x-axis represent Ct values. (**C**) Plaque phenotypes of all SARS-CoV-2 isolate working stocks (BavPat1, IMV1-14) on Vero-CCL81 cells. Monolayers were infected with similar PFUs of each isolate and cultured with an agar overlay for 3 days prior to fixing and staining with crystal violet. Shown is a representative image from each isolate used. (**D**) Quantification of plaque sizes from (C). Individual data points are shown, with means and standard deviations represented. (**E**) Vero-CCL81 cells were infected with the indicated SARS-CoV-2 isolates at an MOI of 0.01 PFU/cell. Supernatants were harvested at several times post-infection and SARS-CoV-2 titers determined by plaque assay. Data represent means and standard deviations from 3 independent experiments, with each experiment performed using technical duplicates. The dotted line crossing the y-axis at 10^2^ PFU/mL indicates the assay limit of detection (LoD). (**F**) Area under the curve (AUC) values for the virus replication data shown in (E), normalized to inoculum titer of the respective isolate. Data represent means and standard deviations from the 3 independent experiments. A one-way ANOVA was performed on log2-transformed data to test for significant differences between the individual isolates (*p*<0.0001). To test for statistical significance against BavPat1, an unpaired t-test was used on log2-transformed data (\**p*<0.05; \*\**p*<0.01; \*\*\**p*<0.001).

Fourteen SARS-CoV-2 primary isolates (IMV1-14) were cultured from samples obtained between 10^th^ March 2020 and 4^th^ May 2020 (highlighted on **Figure 1A**). Together with BavPat1, a SARS-CoV-2 isolate collected on 29^th^ January 2020 in Munich, Germany [30], all viruses were minimally propagated on Vero-CCL81 cells to obtain working stocks (passage 3 (P3) for IMV1-14; passage 2 (P2) for BavPat1). Original Swiss patient material, as well as both P2 and P3 stocks from all isolates, was subjected to next-generation sequencing (NGS) to confirm the absence of additional pathogens, and to provide full-length SARS-CoV-2 genomic sequences (**Supplementary Tables S2 & S3**). Phylogenetic analysis based on the P2 sequences revealed that the isolated viruses were representative of viruses found across Europe during the first half of 2020 [26] (**Supplementary Figure S1**).

As expected, all 14 SARS-CoV-2 isolates harbored several amino acid substitutions when compared to the reference strain SARS-CoV-2/Wuhan-Hu-1, derived from a patient in Wuhan, China in December 2019 [31]. We noted that during working stock preparation (i.e. a total of 3 passages from original patient material in Vero-CCL81 cells), some of the virus isolates acquired additional amino acid substitutions or deletions that were not detected in the original patient material nor in early (P2) passages (**Supplementary Tables S2 & S3**). Most of these passage-induced amino acid substitutions were observed in S, E, or nsps. However, there was no clear consensus change across all isolates that were passaged similarly. The most notable changes were in E, where 6 out of 14 isolates exhibited amino acid substitutions following working stock growth in Vero-CCL81 cells (**Supplementary Table S3**). Furthermore, passage-induced deletion of the furin-cleavage site in S was only observed in 2 out of the 14 isolates (IMV1: Δ677-688, 17%; IMV14: Δ679-685, 81%), and was fully retained in the other 12 isolates. It is possible that the low number of passages that we performed to generate working stocks limited the selection of S furin-cleavage site deletions that have been otherwise readily detected by others [32-34]. We conclude that our set of 14 isolates (IMV1-14) is representative of viruses circulating in Europe in early 2020. Together with their lack of consensus-adaptation to Vero-CCL81 cells during stock preparation, these isolates should therefore be suitable to functionally characterize potential features of early SARS-CoV-2 human adaptation and replication.

### Characterization of SARS-CoV-2 Isolates in Vero-CCL81 Cells and Primary Human Bronchial Epithelial Cells

Using Vero-CCL81 cells, we assessed the plaque phenotypes and growth kinetics of all 14 SARS-CoV-2 isolates, as well as BavPat1. All viruses plaqued well on this substrate, although there was some heterogeneity in plaque size between isolates: for example, as compared to BavPat1, it was striking that IMV4, IMV8, and IMV11 produced smaller plaques (**Figures 1C & D**). Differences between isolates were also observed during multi-cycle growth analysis. While most viruses grew to titers around 10^6^ or 10^7^ PFU/mL over 72h, some viruses were clearly attenuated: final titers of IMV2, IMV8, IMV9, and IMV14 were 10-100-fold lower than the other isolates, and kinetic analysis showed that IMV1, IMV3 and IMV13 grew slower than other isolates (**Figures 1E & F**). Such phenotypic differences were not specific to Vero-CCL81 cells, as we also assessed the replication of a subset of isolates in Vero-E6 cells and noted similar viral growth patterns (**Supplementary Figure S2**).

We next assessed SARS-CoV-2 isolate replication in a previously described primary human bronchial epithelial cell (BEpC) model (**Figure 2A**, [35]). BEpCs were grown at air-liquid interface (ALI) for a minimum of four weeks, and their differentiation into a pseudostratified respiratory epithelium was validated by measuring increased trans-epithelial electrical resistance (TEER) and epithelium-specific cell and tight junction markers, such as β-tubulin and zona occludens protein 1 (ZO-1) (**Supplementary Figures S3A & B**). BEpCs were infected from the apical side with each SARS-CoV-2 isolate and viral titers in both the apical and basolateral compartments were monitored over 72h. Strikingly, there were clear differences in viral replication kinetics between the individual isolates in the apical washes of the infected epithelium (**Figures 2B & C**). BavPat1, as well as 5 other isolates (IMV2, IMV7, IMV9, IMV10, and IMV12) grew to high titers (>10^7^ PFU/mL) within 72 hours. In stark contrast, 2 isolates (IMV6 and IMV4) were unable to replicate in BEpCs at all, while IMV14 was strongly attenuated in this tissue substrate, only reaching titers below 10^4^ PFU/mL. IMV11 also exhibited a notably attenuated growth phenotype, yielding 100-fold lower titers compared to BavPat1, while IMV1, IMV3, IMV5, IMV8, and IMV13 exhibited intermediate growth phenotypes (∼10-fold lower than BavPat1) (**Figures 2B & C**). SARS-CoV-2 titers in the basolateral compartment were minimal as compared to titers in the apical compartment (**Supplementary Figure S3C**), and virus was only ever detectable after 72 hours of infection, suggesting damage of the bronchial epithelium in response to SARS-CoV-2.

**Fig. 2.**
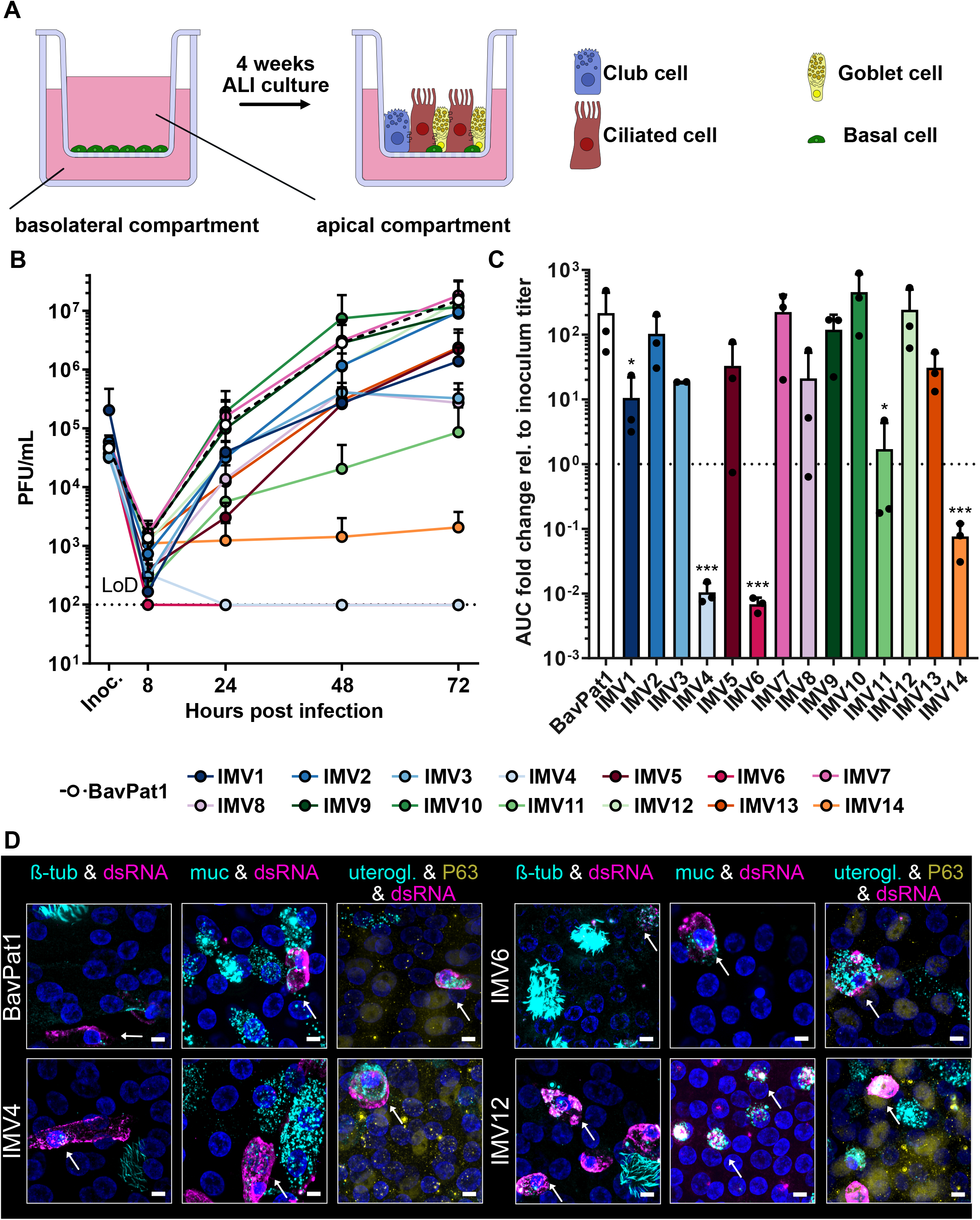
Characterization of SARS-CoV-2 Isolates in Primary Human Bronchial Epithelial Cells. (**A**) Schematic representation of primary human bronchial epithelial cell (BEpC) differentiation into a pseudostratified airway epithelium by culturing at air-liquid interface (ALI) on transwell plates. (**B**) Differentiated BEpCs from donor 1 were infected with 6000 PFU of each SARS-CoV-2 isolate from the apical side. At several times post-infection, apical washes were harvested and virus titers determined by plaque assay. Data represent means and standard deviations from 3 independent experiments, with each experiment performed using 1 individual transwell. The dotted line crossing the y-axis at 10^2^ PFU/mL indicates the assay limit of detection (LoD). (**C**) Area under the curve (AUC) values for the virus replication data shown in (B), normalized to inoculum titer of the respective isolate. Data represent means and standard deviations from the 3 independent experiments. A one-way ANOVA was performed on log2-transformed data to test for significant differences between the individual isolates (*p*<0.0001). To test for statistical significance against BavPat1, an unpaired t-test was used on log2-transformed data (\**p*<0.05; \*\*\**p*<0.001). (**D**) Indirect immunofluorescent staining of SARS-CoV-2 infected BEpCs. Differentiated BEpCs were infected from the apical side with 6000 PFU of the indicated SARS-CoV-2 isolate. At 72 h post-infection, cells were fixed, permeabilized and stained for the presence of infected cells (dsRNA; magenta) and ciliated cells (β-tubulin; cyan), goblet cells (Muc5AC; cyan), club cells (uteroglobin; cyan), or basal cells (P63; yellow). Nuclei were stained with DAPI (blue). Z-stacks were transformed into maximum projection images. Scale bar represents 9.6µm. Arrows indicate co-localization. Data show a subset of representative images from 2 independent experiments performed in different BEpC donors (see also Supplementary Figures S5-10).

We used NGS to fully sequence all viral isolates from BEpC apical compartment samples taken following 72h of replication, but did not observe any mutations leading to amino-acid substitutions as compared to the input Vero-derived P3 stocks used for initial infection. Furthermore, to validate our observations, we generated differentiated BEpCs from two additional human donors and used these new primary cultures to assess the replication capabilities of a subset of virus isolates. While there was some donor-to-donor variation in peak viral titers, most isolates grew very well, but IMV6 and IMV14 were clearly attenuated in all donors, and IMV4 was attenuated in 2 out of 3 donors (**Supplementary Figure S4**).

We next performed indirect immunofluorescent staining in the differentiated human BEpCs to determine the cell tropism of a subset of viral isolates that represented the various growth phenotypes we had previously observed. Using antibodies against β-tubulin, mucin 5AC, uteroglobin, and P63, we identified ciliated, goblet, club, and basal cells, respectively. Viral infection was monitored using an anti-double-stranded (ds) RNA antibody. Despite their reproducible differences in replication capabilities, we found that all isolates displayed a similar cell tropism in this BEpC model of the human respiratory epithelium that was independent of donor source: similar to previous reports [36, 37], we observed SARS-CoV-2 infection of secretory (goblet and club) cells, as well as ciliated cells, although we note that SARS-CoV-2 infected β-tubulin-positive cells often displayed reduced cilia integrity, which has recently been reported by others [38, 39] (**Figure 2D & Supplementary Figures S5-10**). Overall, these SARS-CoV-2 functional data in Vero-CCL81 cells, Vero-E6 cells, and three independent BEpC donors reveal striking differences in replication properties between viral isolates despite a similar cell tropism in primary differentiated human respiratory cells.

### Host-Cell Specific Replication Efficiency of SARS-CoV-2 Isolates

To understand if the spectrum of SARS-CoV-2 isolate replication kinetics was host-cell specific, we compared the relative replication of individual isolates in Vero-CCL81 cells and BEpCs (**Figure 3A**). While some virus isolates replicated very well in both cell types (BavPat1, IMV10, IMV12), other isolates displayed striking host-specific phenotypes. Firstly, IMV2, IMV8 and IMV9 replicated relatively well in BEpCs, but were highly attenuated in Vero-CCL81 cells (**Figure 3A**). These 3 isolates were unique in this phenotype, and each harbored combined amino acid substitutions in Orf3a (Q57H) and nsp2 (T85I) that were not identified in any other isolate (**Supplementary Table S3**), suggestive of an association between these specific changes and the observed cell-specific replication. In contrast, IMV4, IMV6 and IMV11 exhibited a high replication capacity in Vero-CCL81 cells, but were strongly attenuated in BEpCs, with IMV4 and IMV6 being particularly attenuated in most donors (**Figure 3A & Supplementary Figure S4E**). Notably, these 3 isolates each harbored amino acid substitutions at positions 5 (V5G/A) or 6 (S6W) of E (**Supplementary Table S3**). We also noted that IMV14 was highly attenuated in BEpCs and moderately attenuated in Vero-CCL81 cells, which could be due to >80% of the virus population containing a deletion of the furin-cleavage site in S (Δ679-685). As observed by others [24], SARS-CoV-2 isolates such as BavPat1, IMV5, IMV7, IMV10, IMV12 and IMV13 that express S G614 replicated more efficiently in BEpCs than IMV1 and IMV3 that harbor S D614 (**Figure 3A & Supplementary Table S3**). These comparative replication data across 14 different SARS-CoV-2 isolates suggest associations between specific amino acid residues and distinct substrate phenotypes, and imply that the functional viral traits underlying these residues are likely to have important roles for virus replication in human respiratory cells.

**Fig. 3.**
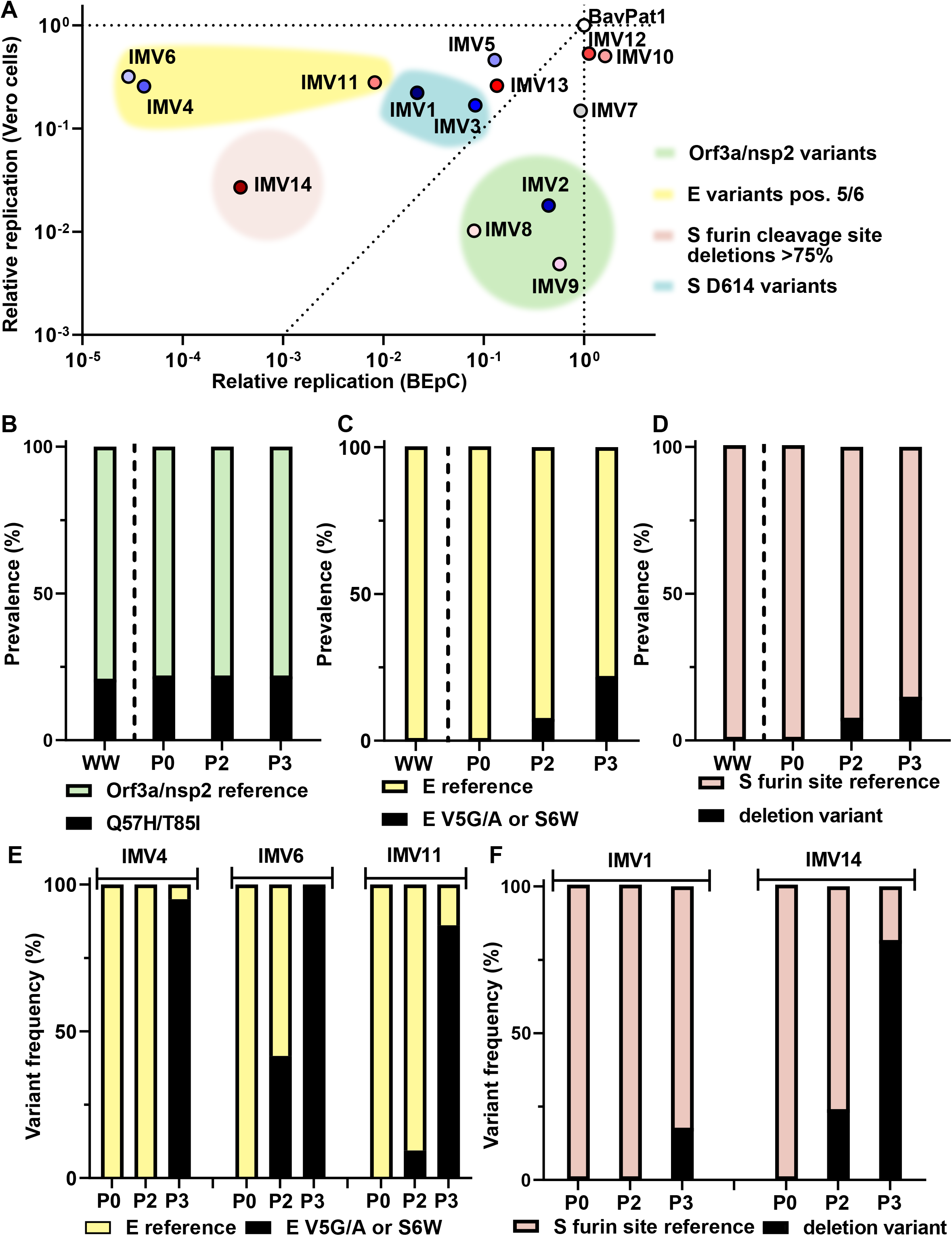
Host-Cell Specific Replication Efficiency of SARS-CoV-2 and Prevalence of Associated Amino Acid Substitutions. (**A**) Relative replication of each SARS-CoV-2 isolate in Vero-CCL81 cells (y-axis, data from Fig. 1F) versus BEpCs (x-axis, data from Fig. 2C) normalized to BavPat1. Groupings of phenotypically similar isolates with common amino acid substitutions are highlighted with colored clouds. (**B-D**) Prevalence of the indicated SARS-CoV-2 amino acid variants in worldwide (WW) published sequences (GISAID, via CoV-GLUE, accessed October 2020), the patient samples (P0), and Vero-CCL81 passages 2/3 (P2/P3) of the SARS-CoV-2 isolates. For worldwide prevalence, published consensus sequences were used, while for patient samples and passages, the available NGS data were analyzed and prevalence was determined based on a frequency of >15%. (**E-F**) Variant frequency of selected SARS-CoV-2 E and S variants within individual patient samples and Vero-CCL81 passages for the indicated isolate was determined by NGS.

### Prevalence of Functional SARS-CoV-2 Amino Acid Substitutions Worldwide and in Laboratory Stocks

We investigated prevalence of the phenotype-associated SARS-CoV-2 isolate variants in Orf3a, nsp2, E, and S in our different Vero-CCL81 passages, original patient material, and worldwide. Orf3a (Q57H) and nsp2 (T85I) variants were found in 20% of our patient samples, a similar prevalence to that found worldwide for the individual variants (**Figure 3B**). Notably, this proportion did not change upon passaging of isolates in Vero-CCL81 cells during stock preparation. In contrast, E protein variants at positions 5/6 and furin-cleavage site deletions in S were all undetectable in worldwide and patient samples, but clearly became more prevalent among our SARS-CoV-2 isolates upon passaging in Vero-CCL81 cells (**Figures 3C & D**). This passaging effect was further exemplified when re-analyzing NGS data from each individual isolate and its parental patient material: the frequency of E S6W, V5G or V5A increased substantially during stock preparation of IMV4, IMV6 and IMV11, respectively (**Figure 3E**), and similar increases in furin-cleavage site deletion variants were observed during stock preparation of IMV1 and IMV14 (**Figure 3F**). These data indicate that, unlike the Orf3a (Q57H) and nsp2 (T85I) variants that exhibit biased replication to BEpCs, the E (V5G/A, S6W) and S furin-cleavage site deletion variants that exhibit biased replication to Vero-CCL81 cells are positively-selected for during passaging in Vero-CCL81 cells.

### Comparative Passaging Reveals Importance of the Spike Furin-Cleavage Site for SARS-CoV-2 Replication in Primary Human Bronchial Epithelial Cells

The furin-cleavage site in SARS-CoV-2 S is highly conserved in patient samples worldwide (**Figure 3D**), and is clearly important for virus pathogenesis [40]. However, a number of studies, including our own here, have reported that deletions in the furin-cleavage site can be acquired during passage in Vero cell substrates [32, 33]. To experimentally assess the importance of the SARS-CoV-2 S furin-cleavage site in primary human respiratory cells, we took advantage of two isogenic IMV14 isolate stocks that exhibited differing frequencies of the S furin-cleavage site deletion (P2, 23.5%; and P3, 81.1%, **Figure 4A**). Comparative passaging for 4 independent replicates in Vero-CCL81 cells and BEpCs revealed that IMV14 P2 (23.5% S furin-cleavage site deletion) replicated in both cell systems (**Figures 4B & 4D**). Furthermore, sequencing of each passage supernatant revealed that, in Vero-CCL81 cells, IMV14 P2 generally increased its frequency of S furin-cleavage site deletion from ∼20% to 50-80% (3 out of 4 replicates within 2 additional passages; **Figure 4C**). In contrast, there was a rapid loss of this deletion variant during passage in BEpCs, with 100% of recovered sequences for all 4 replicates harboring an intact S furin-cleavage site after 2 additional passages (**Figure 4E**). Broadly similar results were obtained when using IMV14 P3 (81.1% S furin-cleavage site deletion): the virus replicated well in Vero-CCL81 cells and retained the S furin-cleavage site deletion at ∼80% frequency (**Figures 4F and 4G**). However, IMV14 P3 was highly attenuated in BEpCs, and virus was only recovered in sufficient quantities from 1 of 4 replicates (**Figure 4H**). Strikingly, sequencing of this BEpC-recovered virus revealed a rapid loss of the S furin-cleavage site deletion (**Figure 4I**). Together with work from others, these data provide strong experimental evidence that the S furin-cleavage site is an essential viral trait required, and selected, for efficient SARS-CoV-2 replication in primary human respiratory cells [40, 41].

**Fig. 4.**
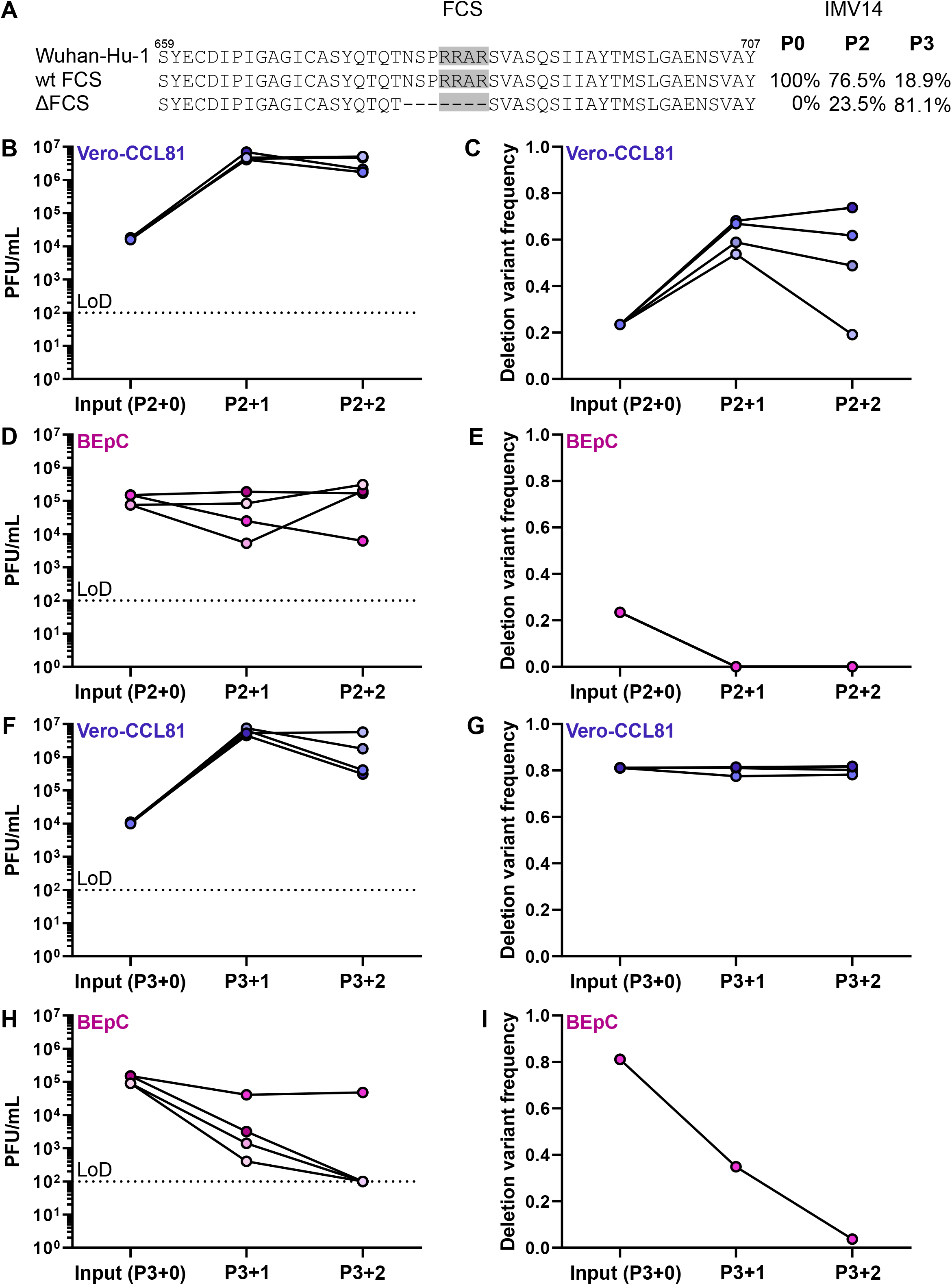
The SARS-CoV-2 Spike Furin-Cleavage Site is Essential for Efficient Replication in Primary Human Bronchial Epithelial Cells. (**A**) Amino acid sequences surrounding the Spike (S) furin-cleavage site (FCS, gray) in SARS-CoV-2 isolate IMV14, as compared to the reference sequence, Wuhan-Hu-1 (MN908947.3). Percentages on the right show the frequency of the indicated variant in IMV14 P2 and IMV14 P3 stocks that were used in subsequent experiments. (**B-E**) IMV14 P2 stock (23.5% S furin-cleavage site deletion frequency) was passaged twice on Vero-CCL81 cells or BEpCs. At each passage, virus titers were determined by plaque assay (B & D) and frequency of the furin-cleavage site deletion was determined by NGS (C & E). Data represent the individual results from 4 independent replicates. (**F-I**) IMV14 P3 stock (81.1% S furin-cleavage site deletion frequency) was passaged twice on Vero-CCL81 cells or BEpCs. At each passage, virus titers were determined by plaque assay (F & H) and frequency of the furin-cleavage site deletion was determined by NGS (G & I). Data represent the individual results from 4 independent replicates. For E and I, a sequence could only be determined from 1 replicate, where limited SARS-CoV-2 replication occurred. In panels B, D, F & H, the dotted lines crossing the y-axes at 10^2^ PFU/mL indicate the assay limit of detection (LoD).

## DISCUSSION

Herein, our functional characterization of a spectrum of first wave SARS-CoV-2 isolates has revealed several viral traits that are critical, and potentially adaptive, for efficient viral replication in human respiratory cells:

Firstly, using fully replication-competent SARS-CoV-2 isolates in primary human bronchial epithelial cells, we observe that viruses expressing the naturally-occurring G614 substitution in S generally replicate more efficiently than otherwise similar viruses expressing the parental D614 version of S. These observations are in-line with previous work using virus-like particle or vesicular stomatitis virus (VSV) pseudotype systems, where S G614 pseudotypes were found to be more infectious than S D614 pseudotypes [22, 23]. Furthermore, recent SARS-CoV-2 reverse genetics studies have also linked the single D614G substitution to increased infectivity and virus replication in primary human cells as well as *in vivo* models [24, 25]. Mechanistically, the D614G substitution appears to permit a more open ACE2-binding conformation of the S trimer, which may lead to increased virus fusion with host cell membranes [22]. Together with our results from naturally-occurring SARS-CoV-2 isolates, these common findings using different experimental systems provide a clear basis to understand the rapid emergence and population-wide spread of the S D614G substitution.

Secondly, we uncovered an association between naturally-occurring Orf3a (Q57H) and nsp2 (T85I) variants, and a cell-specific replication phenotype: SARS-CoV-2 isolates unique in harboring these variants replicated poorly in Vero-CCL81 cells, while maintaining efficient replicative capacity in primary human bronchial epithelial cells. These variants are found in approximately 20% of SARS-CoV-2 sequences worldwide, but their functional consequences are unclear. The functions of SARS-CoV-2 nsp2 have not been fully elucidated, although it has been described to interact with host proteins involved in endomembrane compartments, vesicle trafficking and translation [42]. Orf3a is an ion channel with pro-apoptotic and pro-inflammatory functions [42-44]. Recent structural studies have revealed that Q57 forms the major hydrophilic constriction in the SARS-CoV-2 Orf3a pore, however, the Q57H substitution does not appear to influence Orf3a channel activities [44]. Nevertheless, the Orf3a Q57H variant was acquired shortly after introduction into humans, and is not found in early SARS-CoV-2 sequences or in related bat coronaviruses [44], potentially suggesting positive-selection of this residue similar to S D614G. Notably, the mutation leading to the Orf3a Q57H variant also changes the sequence of two other putative (and poorly characterized) overlapping reading frame products, Orf3c and Orf3d [45, 46]. Further studies, including reverse genetics experiments, will be required to determine whether these variants are functionally linked to one another, and how each variant may mechanistically determine this phenotype. While Vero-CCL81 cells are clearly not a physiological substrate for SARS-CoV-2, it may be that the cell-specific replication capacities that we observe with isolates expressing Orf3a (Q57H) and nsp2 (T85I) variants also occurs in other, non-respiratory, cell-types of the human body, potentially impacting disease pathogenesis.

Thirdly, we observed that SARS-CoV-2 stock preparation in Vero-CCL81 cells led many of the isolates (6 out of 14) to acquire passage-derived amino acid substitutions in the E protein, which were not observed in patient material (either from our samples or worldwide). Strikingly, substitutions at E positions 5/6 (V5G/A or S6W) were highly attenuated in primary human bronchial epithelial cells but replicated well, and were potentially selected for, in Vero-CCL81 cells. Previous work using SARS-CoV-1 has shown that E deletion viruses are only mildly attenuated in Vero-E6 cells as compared to their attenuation in human cell-lines or in *in vivo* models [16], and such viruses have been considered as the basis for live-attenuated vaccine concepts [47-50]. These data, together with the observations presented here, suggest that SARS-CoV-2 E has important species- or cell-type-specific functions, and further indicate that residues at positions 5 and 6 play key roles in some critical E activities. It will be interesting in future studies to determine whether such activities include slowing of the cell secretory pathway by E [51], and/or the activation of host inflammasomes by E [52], as well as how the requirement for these functions differs between species or cell-types.

Finally, as also observed by others [32-34, 53], we found that in-frame deletions of the furin-cleavage site in S could be selected for and enriched in SARS-CoV-2 isolates during passage in Vero-CCL81 cells. This was apparent even for very low passage stocks as described here (passage 3 from original patient material), but importantly was not a universal feature of all SARS-CoV-2 isolates: deletions of the S furin-cleavage site were only detected in 2 out of 14 of our primary isolates. The furin-cleavage site permits ‘priming’ of S (cleavage at S1/S2 boundary) by furin-like proteases [11], thereby enabling the virus to fuse at the cellular plasma membrane if TMPRSS2 is present to further cleave S at the S2’ site [12, 13]. In the absence of furin-cleavage, ‘priming’ and subsequent cleavage of S at the S2’ site occurs in endosomes by cathepsin proteases. Vero cells appear to lack expression of TMPRSS2 [13, 33], and thus maintaining the S furin-cleavage site is unlikely to be advantageous to SARS-CoV-2 in these cells. This provides a plausible mechanistic basis for SARS-CoV-2 adaptation to Vero cells through deletions in the S furin-cleavage site. Such findings, together with our additional observations relating to functional E substitutions during SARS-CoV-2 growth in Vero-CCL81 cells, underscore the need for care during stock preparation, particularly where viruses are to be compared with one another for phenotypes. Thus, it may be that other cell-lines, including human Calu-3 [53], are more suitable for SARS-CoV-2 propagation, and should be thoroughly characterized in the future as a possible substrate to limit cell culture adaptation in SARS-CoV-2 genes such as S and E. Nevertheless, we found that in-frame deletions in the S furin-cleavage site attenuated SARS-CoV-2, most notably in primary human bronchial epithelial cells. However, passaging of SARS-CoV-2 preparations containing the S furin-cleavage site deletions in primary human bronchial epithelial cells provided a very high selection pressure to enrich only for viruses that retained the full S furin-cleavage site. Thus, our study of SARS-CoV-2 cell adaptations revealed the critical nature of this S furin-cleavage site for SARS-CoV-2 replication in primary human respiratory cells, a finding similar to that recently described by others using alternative methods [40, 41]).

In sum, our work identifies both key features in the SARS-CoV-2 genome that are essential for its growth in human respiratory cells, as well as variants derived during human circulation that determine efficient replication.

## MATERIALS & METHODS

### Cells

Vero-CCL81 cells (ATCC) and Vero-E6 cells (kindly provided by Volker Thiel, University of Bern, Switzerland) were cultured at 37°C and 5% CO_2_ in Dulbecco’s Modified Eagle’s Medium (DMEM; Gibco), supplemented with 10% (v/v) fetal calf serum (FCS), 100 U/mL of penicillin and 100 μg/mL of streptomycin (#15140-122; Gibco). Primary human bronchial epithelial cells (BEpCs) from a 73 year-old female donor (donor 1) were purchased from Promocell (#C-12640). BEpCs from a 53 year-old male donor (donor 2) and a 62 year-old male donor (donor 3) were purchased from Epithelix (#EP51AB). Cells were grown in airway epithelium basal growth medium (Promocell, #C-21260) supplemented with an airway growth medium supplement pack (#C-39160; Promocell) and 10 µM Y-27632 (Selleck Chemicals). Differentiation of BEpCs, and subsequent validation of airway cultures by measuring the transepithelial electrical resistance (TEER) and immunofluorescence, was performed exactly as recently described [35].

### SARS-CoV-2 Isolation and Stocks

SARS-CoV-2 isolates (termed IMV1-14: SARS-CoV-2/human/Switzerland/ZH-UZH-IMV1/2020 - SARS-CoV-2/human/Switzerland/ZH-UZH-IMV14/2020 following ICTV guidelines [54], GISAID [55] accession IDs EPI_ISL_590823 to EPI_ISL_590836) were isolated on Vero-CCL81 cells essentially as described [35, 56], using anonymized patient samples that had tested PCR-positive during routine diagnostics at the Institute of Medical Virology, University of Zurich between March and May 2020. Briefly, 100 μL of a 5-fold dilution series of sample in serum-free DMEM was mixed with 1×10^5^ Vero-CCL81 cells in 500 µL of DMEM supplemented with 10% FCS, 100 U/mL penicillin, 100 μg/mL streptomycin, and 2.5 µg/mL amphotericin B (#15290018; Gibco). Cells were seeded in 24-well plates and incubated at 37°C for 4-5 days until cytopathic effect was apparent. Supernatants were centrifuged at 1500 rpm for 5 minutes, and 250 μL of this cleared supernatant (termed passage 1; P1) was used to inoculate a 25 cm^2^ flask of freshly seeded Vero-CCL81 cells, which were then cultured for a further 4-5 days at 37°C. Cell supernatants were harvested, clarified by centrifugation at 1500 rpm for 5 minutes, and aliquoted before freezing at -80°C (termed P2). P2 stocks were verified by our in-house diagnostics service to be PCR-positive for SARS-CoV-2. Following titer determination by plaque assay [35], a P3 working stock was generated by infecting Vero-CCL81 cells at an MOI of 0.001 PFU/cell for 72h in DMEM supplemented with 100 U/mL penicillin, 100 μg/mL streptomycin, 0.3% bovine serum albumin (BSA; (#A7906, Sigma-Aldrich), 20 mM HEPES (#H7523, Sigma-Aldrich), 0.1% FCS, and 0.5 μg/mL TPCK-treated trypsin (#T1426, Sigma-Aldrich), prior to supernatant clarification by centrifugation (1500 rpm, 5 mins), aliquoting/storage at -80°C, and plaque titration. BetaCoV/Germany/BavPat1/2020 was obtained from the European Virus Archive GLOBAL (EVA-GLOBAL; Ref-SKU: 026V-03883) [30]. All work with infectious SARS-CoV-2 was performed in an approved BSL3 facility by trained personnel at the Institute of Medical Virology, University of Zurich. All procedures and protective measures were thoroughly risk assessed prior to starting the project, and were approved by the Swiss Federal Office of Public Health (Ecogen number A202808/3).

### SARS-CoV-2 Full Genome Sequencing

SARS-CoV-2 genome sequencing from P2 and P3 stocks was performed using a next-generation sequencing (NGS) approach as described previously [57]. Briefly, samples were diluted in NucliSENS EasyMAG lysis buffer (BioMérieux, Craponne, France; ratio 1:4) and filtered using a 0.45 µm PES filter (TPP, Trasadingen, Switzerland). Total nucleic acids were extracted using the NucliSENS EasyMAG system, followed by reverse transcription with random hexamers and second strand synthesis. Sequencing libraries were constructed using the NexteraXT protocol (Illumina, San Diego, CA, USA) and sequenced on an Illumina MiSeq for 1 x 151 cycles using version 3 chemistry.

SARS-CoV-2 genome sequencing from original patient material was performed using a previously described targeted, tiled-amplicon, NGS approach [58, 59]. Briefly, total nucleic acids were extracted followed by reverse transcription with random hexamers and oligo-dT priming (ratio 3:1) using SuperScript IV Reverse Transcriptase (Thermo Fisher Scientific). The generated cDNA was used as input for 14 overlapping PCR reactions (ca. 2.5□Lkb each) spanning the entire viral genome using Platinum SuperFi DNA Polymerase (Thermo Fisher Scientific). Amplicons were pooled per patient before NexteraXT library preparation and sequencing on an Illumina MiSeq for 1 x 151 cycles.

### SARS-CoV-2 Genome Analysis

To generate SARS-CoV-2 consensus sequences, all reads were iteratively aligned using SmaltAlign (github.com/medvir/SmaltAlign). Variant calling was performed as follows: raw sequences were mapped to a reference sequence (MN985325.1) using minimap2 (version 2.17-r941) [60] and variants (≥ 15% frequency). were called using lofreq (version 2.1.5) [61]. The phylogenetic analysis by maximum likelihood was performed on the 14 sequences of P2 and a set of 569 representative SARS-CoV-2 sequences used by Alm et al. [26] downloaded from GISAID [55]. The sequences were aligned using MAFFT v7.271 [60] and the phylogeny was estimated using RAxML [61] with GTR+G+F model. Consensus sequences were analyzed using CoV-GLUE [62] to identify amino acid variants and SNPs, as well as to assign viral lineages according to nomenclature proposals [63]. Clades were determined by nextstrain.org [64].

### SARS-CoV-2 Replication Assays

For Vero-CCL81 and Vero-E6 experiments, 1.2×10^4^ cells were seeded overnight in 96-well plates. Cells were then infected with SARS-CoV-2 at an MOI of 0.01 PFU/cell in PBS supplemented with 0.3% BSA, 1 mM Ca^2+^/Mg^2+^, 100 U/mL penicillin, and 100 μg/mL streptomycin. After 1h inoculation, cells were washed once in PBS, and the medium was replaced with DMEM supplemented with 100 U/mL penicillin, 100 μg/mL streptomycin, 0.3% BSA, 20 mM HEPES, 0.1% FCS, and 0.5 μg/mL TPCK-treated trypsin. Samples were taken at the indicated time points and stored at -80°C prior to titer determination by plaque assay [35]. Plaque size quantification was performed using ImageJ analysis software. For BEpC experiments, cells grown on 6.5 mm transwell filter inserts were infected from the apical side with 6000 PFU of the respective SARS-CoV-2 isolate in PBS supplemented with 0.3% BSA, 1 mM Ca^2+^/Mg^2+^, 100 U/mL penicillin, and 100 μg/mL streptomycin. After 1h, the inoculum was removed and cells were incubated at 37°C. At selected time points, virus was harvested from the apical side by applying 80 µL of PBS for 15 min at 37°C before removal and storage at -80°C prior to titer determination by plaque assay [35]. Basolateral samples were also collected and frozen at -80°C.

### SARS-CoV-2 Passaging Assays

For Vero-CCL81 experiments, ∼2×10^5^ cells were infected with the indicated SARS-CoV-2 isolate stock at an MOI of 0.01 PFU/cell as described above. In parallel, BEpCs grown on a 12mm filter insert were infected from the apical side with 2.2×10^4^ PFU/well. After 72h at 37°C, virus was harvested by either directly collecting the supernatant from infected Vero-CCL81 cells, or by performing apical washes of BEpCs with PBS. One third of the harvested supernatant (passage 1; P1) was used to similarly infect fresh Vero-CCL81 cells or BEpCs for a subsequent passage (P2), while the remaining supernatant was used for full-genome sequencing (see above) and SARS-CoV-2 titration [35].

### Immunofluorescence

Cells were fixed with 3.7% paraformaldehyde in PBS and permeabilized with PBS supplemented with 50 mM ammonium chloride (#254134; Sigma-Aldrich), 0.1% saponin (#47036, Sigma-Aldrich) and 2% BSA (#A7906; Sigma-Aldrich). A mouse (#Ab00458-1.1) or rabbit (Ab00458-23.0) anti-dsRNA antibody (9D5; Lucerna-Chem) was used to stain for SARS-CoV-2 infected cells. A mouse anti-β-tubulin IV antibody (#ab11315; Abcam), a mouse anti-MUC5AC antibody (#ab3649; Abcam), a rabbit anti-P63 antibody (#ab124762; Abcam), and a rat anti-uteroglobin antibody (#MAB4218; R&D Systems) was used to stain ciliated, goblet, basal, and club cells, respectively. As secondary antibodies, anti-mouse IgG Alexa488 (#A-11029), anti-mouse IgG Alexa647 (#A-31571), anti-rabbit IgG Alexa546 (#A-10040), anti-rabbit IgG Alexa647 (#A-31573), and anti-rat IgG Alexa488 (#A-11006) antibodies were used (all from Thermo Fisher Scientific). Nuclei were stained with DAPI (#10236276001; Sigma Aldrich). Filters were mounted using ProLong Gold Antifade Mountant (#P36930; Thermo Fisher Scientific) and z-stack images were acquired using a DMi8 microscope (Leica) and processed using the THUNDER Large Volume Computational Clearing algorithm (Leica). Maximum projection images of z-stacks were generating using ImageJ software.

### Statistical Analyses

Statistical significance was determined using one-way ANOVA and unpaired t-test on log2-transformed data.

## ACKNOWLEDGEMENTS

We thank members of the Institute of Medical Virology Diagnostics service, as well as Peter Rusert, Jacqueline Weber, and Sonja Fernbach (University of Zurich, UZH), for excellent technical assistance. Vero-E6 cells were kindly provided by Volker Thiel (University of Bern, Switzerland). Imaging was performed with equipment maintained by the Center for Microscopy and Image Analysis, UZH. We gratefully acknowledge originating and submitting laboratories where SARS-CoV-2 genetic sequence data were generated and shared via the GISAID initiative. This work was partially supported by the UZH through core Institute funds, a Pandemiefonds grant of the UZH Foundation (to AT), a UZH Forschungskredit grant (FK-18-044 to IB), and the UZH Clinical Research Priority Program ‘Comprehensive Genomic Pathogen Detection’ (to MH and AT). The senior authors’ laboratories are funded by the Swiss National Science Foundation through grants 31003A_182464 to BGH and 31003A_176170 to SSt.

## FIGURE LEGENDS

**Supplementary Fig. S1. Phylogenetic Analysis of the SARS-CoV-2 Isolates Generated in this Study**. Phylogenetic analysis of sequences derived from the 14 SARS-CoV-2 P2 isolates, together with the SARS-CoV-2 sequences from Alm *et al*. [26] that represent viral diversity across the WHO European Region during the same timeframe. Wuhan/WH04/2020 (EPI_ISL_406801), belonging to clade 19B, was chosen as the outgroup. The tree is colored based on SARS-CoV-2 Nextstrain clades, with circles indicating the positions of 14 SARS-CoV-2 isolates generated in this study. The scale bar indicates the number of nucleotide substitutions per site.

**Supplementary Fig. S2. Characterization of Selected SARS-CoV-2 Isolates in Vero-E6 Cells**. (**A**) Vero-E6 cells were infected with the indicated SARS-CoV-2 isolates at an MOI of 0.01 PFU/cell. Supernatants were harvested at the indicated times post-infection and SARS-CoV-2 titers determined by plaque assay. Data represent means and standard deviations from 3 independent experiments, with each experiment performed using 1 individual well. The dotted line crossing the y-axis at 10^2^ PFU/mL indicates the assay limit of detection (LoD). (**B**) Area under the curve (AUC) values for the virus replication data shown in (A), normalized to inoculum titer of the respective isolate. Data represent means and standard deviations from the 3 independent experiments. A one-way ANOVA was performed on log2-transformed data to test for significant differences between the individual isolates (*p*<0.0001). To test for statistical significance against BavPat1 an unpaired t-test was used on log2-transformed data (\*\**p*<0.005; \*\*\**p*<0.0005).

**Supplementary Fig. S3. Characterization and Validation of Primary Human Bronchial Epithelial Cell Cultures**. (**A**) During primary human bronchial epithelial cell (BEpC, donor 1) differentiation into a pseudostratified airway epithelium at the air-liquid interface (ALI), transepithelial electrical resistance (TEER) was measured weekly. Data represent mean TEER values and standard deviations from 3 independent wells at each time-point. (**B**) Differentiated BEpCs from donor 1 were fixed and stained for ciliated cells (β-tubulin; cyan), tight junctions (ZO-1; magenta) and nuclei (DAPI; blue). Scale bar represents 25µm. (**C**) Differentiated BEpCs from donor 1 were infected from the apical side with 6000 PFU of each SARS-CoV-2 isolate (see Figure 2B). At the indicated times post-infection, basolateral samples were harvested and virus titers were determined by plaque assay. Data represent means and standard deviations from 2 independent replicates. The dotted line crossing the y-axis at 10^2^ PFU/mL indicates the assay limit of detection (LoD).

**Supplementary Fig. S4. Characterization of Selected SARS-CoV-2 Isolates in Additional Primary Human Bronchial Epithelial Cell Donors**. (**A & C**) Differentiated

BEpCs from donors 2 (A) and 3 (C) were infected with 6000 PFU of the indicated SARS-CoV-2 isolate from the apical side. At the indicated times post-infection, apical washes were harvested and virus titers were determined by plaque assay. Data represent means and standard deviations from 3 independent replicates. The dotted lines crossing the y-axes at 10^2^ PFU/mL indicate the assay limit of detection (LoD). (**B & D**) Area under the curve (AUC) values for the virus replication data shown in (A & C), normalized to inoculum titer of the respective isolate. Data represent means and standard deviations from the 3 independent experiments. A one-way ANOVA was performed on log2-transformed data to test for significant differences between the individual isolates (n.s. for donor 2; *p*<0.0001 for donor 3). To test for statistical significance against BavPat1 (D) an unpaired t-test was used on log2-transformed data (\**p*<0.05; \*\**p*<0.01). Of note, as BavPat1 exhibited an unexplained attenuated phenotype in BEpCs from donor 2 (B), an unpaired t-test was used on log2-transformed data to test for statistical significance against IMV3 (\**p*<0.05; \*\**p*<0.01). (**E**) Boxplot representation of the area under the curve (AUC) values for the virus replication data from the 3 independent donors (data from B & D; and Fig. 2C), normalized to inoculum titer of the respective isolate and donor. Shown is median and standard deviation. The red triangles represent data obtained from donor 1.

**Supplementary Fig. S5. Indirect Immunofluorescent Staining of SARS-CoV-2 Infected Cells and Ciliated Cells in BEpCs from Donor 2**. Differentiated BEpCs from donor 2 were infected from the apical side with 6000 PFU of the indicated SARS-CoV-2 isolate. At 72 h post-infection cells were fixed, permeabilized and stained for the presence of infected cells (dsRNA; magenta) and ciliated cells (β-tubulin; cyan). Nuclei were stained with DAPI (blue). Arrows indicate co-localization. Scale bars represent 50µm for the 20x magnifications, and 8µm for the 100x magnifications. Representative maximum projection images of z-stacks from one experiment are shown.

**Supplementary Fig. S6. Indirect Immunofluorescent Staining of SARS-CoV-2 Infected Cells and Goblet Cells in BEpCs from Donor 2**. Differentiated BEpCs from donor 2 were infected from the apical side with 6000 PFU of the indicated SARS-CoV-2 isolate. At 72 h post-infection cells were fixed, permeabilized and stained for the presence of infected cells (dsRNA; magenta) and goblet cells (MUC5AC; cyan). Nuclei were stained with DAPI (blue). Arrows indicate co-localization. Scale bars represent 50µm for the 20x magnifications and 8µm for the 100x magnifications. Representative maximum projection images of z-stacks from one experiment are shown.

**Supplementary Fig. S7. Indirect Immunofluorescent Staining of SARS-CoV-2 Infected Cells, Club Cells, and Basal Cells in BEpCs from Donor 2**. Differentiated BEpCs from donor 2 were infected from the apical side with 6000 PFU of the indicated SARS-CoV-2 isolate. At 72 h post-infection cells were fixed, permeabilized and stained for the presence of infected cells (dsRNA; magenta), club cells (uteroglobin; cyan), and basal cells (P63; yellow). Nuclei were stained with DAPI (blue). Arrows indicate co-localization. Scale bars represent 50µm for the 20x magnifications and 8µm for the 100x magnifications. Representative maximum projection images of z-stacks from one experiment are shown.

**Supplementary Fig. S8. Indirect Immunofluorescent Staining of SARS-CoV-2 Infected Cells and Ciliated Cells in BEpCs from Donor 3**. Differentiated BEpCs from donor 3 were infected from the apical side with 6000 PFU of the indicated SARS-CoV-2 isolate. At 72 h post-infection cells were fixed, permeabilized and stained for the presence of infected cells (dsRNA; magenta) and ciliated cells (β-tubulin; cyan). Nuclei were stained with DAPI (blue). Arrows indicate co-localization. Scale bars represent 50µm for the 20x magnifications, and 8µm for the 100x magnifications. Representative maximum projection images of z-stacks from one experiment are shown.

**Supplementary Fig. S9. Indirect Immunofluorescent Staining of SARS-CoV-2 Infected Cells and Goblet Cells in BEpCs from Donor 3**. Differentiated BEpCs from donor 3 were infected from the apical side with 6000 PFU of the indicated SARS-CoV-2 isolate. At 72 h post-infection cells were fixed, permeabilized and stained for the presence of infected cells (dsRNA; magenta) and goblet cells (MUC5AC; cyan). Nuclei were stained with DAPI (blue). Arrows indicate co-localization. Scale bars represent 50µm for the 20x magnifications and 8µm for the 100x magnifications. Representative maximum projection images of z-stacks from one experiment are shown.

**Supplementary Fig. S10. Indirect Immunofluorescent Staining of SARS-CoV-2 Infected Cells, Club Cells, and Basal Cells in BEpCs from Donor 3**. Differentiated BEpCs from donor 3 were infected from the apical side with 6000 PFU of the indicated SARS-CoV-2 isolate. At 72 h post-infection cells were fixed, permeabilized and stained for the presence of infected cells (dsRNA; magenta), club cells (uteroglobin; cyan), and basal cells (P63; yellow). Nuclei were stained with DAPI (blue). Arrows indicate co-localization. Scale bars represent 50µm for the 20x magnifications and 8µm for the 100x magnifications. Representative maximum projection images of z-stacks from one experiment are shown.

