## Supplemental Table 1 for "SARS-CoV-2 Variants Reveal Features Critical for Replication in Primary Human Cells"

| Sample number | Ct custom PCR (E) | Ct Cobas (ORF1/E) | Success/ Failure | Sample number | Ct custom PCR (E) | Ct Cobas / ORF1/E) | Success/ Failure |
|---|---|---|---|---|---|---|---|
| 1 | 17.86 | - | success | 34 | - | 30.91/33.24 | failure |
| 2 | 19.4 | - | success | 35 | - | 32.35/33.52 | failure |
| 3 | 18.95 | - | failure | 36 | - | 32.58/34.62 | failure |
| 4 | 19.98 | - | success | 37 | - | 32.34/33.37 | failure |
| 5 | 18.56 | - | failure | 38 | - | 30.97/31.44 | failure |
| 6 | 19.5 | - | failure | 39 | - | 32.21/33.62 | failure |
| 7 | 19.73 | - | failure | 40 | - | 33.33/34.8 | failure |
| 8 | 19.34 | - | failure | 41 | - | 30.88/32.18 | failure |
| 9 | 19.4 | - | success | 42 | - | 31.82/33.3 | failure |
| 10 | 17 | - | success | 43 | - | neg/37.85 | failure |
| 11 | 18.5 | - | success | 44 | - | 34.93/36.44 | failure |
| 12 | - | 19.92/19.98 | failure | 45 | 37.42 | - | failure |
| 13 | 19.4 | - | success | 46 | - | 35.09/neg | failure |
| 14 | - | 17.97/18.22 | success | 47 | - | 33.79/37.08 | failure |
| 15 | 21.75 | - | success | 48 | - | 35.23/38.12 | failure |
| 16 | 21 | - | success | 49 | - | neg/38.71 | failure |
| 17 | - | 21.67/21.46 | success | 50 | - | 33.67/35.09 | failure |
| 18 | - | 21.35/21.5 | failure | 51 | - | 33.66/35.86 | failure |
| 19 | - | 20.04/20.16 | success | 52 | - | 35.12/36.64 | failure |
| 20 | - | 20.05/20.22 | success | 53 | - | 34.31/37.74 | failure |
| 21 | - | 24.81/24.47 | failure | 54 | - | 32.6/35.16 | failure |
| 22 | - | 28.73/29 | failure | 55 | - | 34.5/36.28 | failure |
| 23 | - | 28.96/29.48 | success | 56 | - | 34.44/37.54 | failure |
| 24 | - | 28.58/29.16 | failure | 57 | - | 34.12/36.14 | failure |
| 25 | - | 28.48/29.23 | failure | 58 | - | neg/36.95 | failure |
| 26 | - | 29.67/29.91 | failure | 59 | - | neg/37.96 | failure |
| 27 | - | 26.55/26.82 | failure | 60 | - | neg/37.25 | failure |
| 28 | - | 28.8/28.92 | failure | 61 | - | 34.71/35.97 | failure |
| 29 | - | 27.7/27.96 | failure | 62 | - | neg/36.68 | failure |
| 30 | - | 27.39/27.91 | failure | 63 | - | 33.79/37.08 | failure |
| 31 | - | 25.39/25.64 | failure | 64 | - | 33.67/35.09 | failure |
| 32 | - | 31.97/34.11 | failure | 65 | - | 33.66/35.86 | failure |
| 33 | - | 30.27/31.49 | failure | 66 | - | 32.6/35.16 | failure |
| | | | | 67 | - | 33.28/35.2 | failure |

**Supplementary Table S1. Cycle threshold (Ct) values, and success/failure outcomes of virus isolation attempts, from 67 patient samples that were PCR-positive for SARS-CoV-2**
