## Supplemental Table 2 for "SARS-CoV-2 Variants Reveal Features Critical for Replication in Primary Human Cells"

| **ID (Date)*^a^*** | **Lineage or**  **Clade*^b^*** | **Amino acid variants*^c^*** | | | | | | | **SNPs*^c^*** |
| --- | --- | --- | --- | --- | --- | --- | --- | --- | --- |
|  |  | **Non-structural proteins** | **S** | **Orf3a** | **E** | **M** | **Orf7a** | **N** |  |
| **IMV1**  (10/03) | B.2.2 / 19A | nsp1 (V111V/L)*^d^*  nsp2 (H237R)  **nsp2 (D582D/A)***^d^*  **nsp6 (L37F)** | G72R/G*^d^*  D215D/G*^d^* | **G251V** |  |  |  |  | G596T, A1515G, A2550C, C9223T, C11074T, G11083T, C14805T, T17247C, G21776A, A22206G, G26144T |
| **IMV2**  (09/03) | B.1 / 20C | **nsp2 (T85I)**  **nsp12 (P323L)** | **D614G** | **Q57H***^e^* |  |  |  |  | C1059T, C3037T, C14408T, A23403G, G25563T |
| **IMV3**  (11/03) | B.2 / 19A | nsp2 (P624P/L)*^d^*  **nsp6 (L37F)** |  | **G251V** |  |  |  |  | C2676T, G11083T, C14805T, T17247C, C25626T, G26144T |
| **IMV4**  (11/03) | B.1 / 20A | **nsp12 (P323L)** | **D614G** |  |  |  |  |  | C3037T, C14408T, C15324T, A23403G |
| **IMV5**  (11/03) | B.1.1 / 20B | **nsp12 (P323L)** | **D614G** |  |  | **T175M** |  | **R203K**  **G204R** | C3037T, C14408T, A23403G, C27046T, G28881A, G28882A, G28883C |
| **IMV6**  (29/03) | B.1.5 / 20A | **nsp3 (K1693N)**  **nsp12 (P323L)** | **D614G** |  | V5G/V*^d^* |  |  | **S194L***^e^* | C3037T, G7798T, C14408T, A20268G, A23403G, T26258G, C28854T |
| **IMV7**  (29/03) | B.1 / 20A | nsp3 (H795H/Y)*^d^*  nsp3 (S915N/S)*^d^*  nsp12 (P323L) | D614G |  |  |  |  |  | C2638A, C3037T, C5102T, G5463A, C14408T, C15324T, C21691T, A23403G |
| **IMV8**  (30/03) | B.1 / 20C | **nsp2 (T85I)**  **nsp12 (P323L)** | **S514F**  **D614G** | **Q57H***^e^* |  |  |  |  | C1059T, C3037T, C14408T, C23103T, A23403G, G25563T, C29535T |
| **IMV9**  (07/04) | B.1 / 20C | **nsp2 (T85I)**  nsp4 (N405K/N)*^d^*  nsp12 (A97A/V)*^d^*  **nsp12 (P323L)**  **nsp12 (S647I)**  nsp15 (S312T/S)*^d^*  nsp15 (S315S/F)*^d^* | Q218K/Q*^d^*  **D614G** | **Q57H***^e^* |  |  |  |  | C1059T, C3037T, G7459A, T9769A, C13730T, C14408T, G15380T, T20554A, C20564T, C22214A, A23403G, G25563T |
| **IMV10**  (14/04) | B.1.1 / 20B | **nsp4 (S336L)**  **nsp12 (P323L)** | **D614G** |  |  |  |  | **R203K**  **G204R**  **I292T** | C2462T, C3037T, C9561T, C14408T, T15837C, A23403G, G28881A, G28882A, G28883C, T29148C |
| **IMV11**  (04/05) | B.1.5 / 20A | **nsp3 (T970M)**  **nsp12 (P323L)** | **E583D**  **D614G** |  |  |  |  | **S194L***^e^* | C3037T, C5628T, A9529T, C14408T, A20268G, G23311T, A23403G, C28854T |
| **IMV12**  (04/05) | B.1.1 / 20B | **nsp12 (P323L)** | **D614G** |  |  |  |  | **R203K**  **G204R** | C3037T, T11449C, C14408T, A20979G, A23403G, G28881A, G28882A, G28883C |
| **IMV13**  (04/05) | B.1 / 20A | **nsp6 (L37F)**  **nsp12 (P323L)** | **D614G** |  |  |  | **H47N** | R203K/R*^d^* | C3037T, G11083T, C14408T, C15324T, A23403G, C27532A, G28881A |
| **IMV14**  (04/05) | B.1 / 20A | **nsp2 (I273L)**  nsp3 (K1083K/R)*^d^*  **nsp8 (A3V)**  **nsp12 (P323L)**  nsp16 (D108D/A)*^d^* | **D614G**  Δ679-685*^d^* |  |  |  |  | **L139F** | C556T, A1622C, C3037T, A5967G, C12099T, C14408T, C15324T, A20981C, A23403G, G28690T |

**Supplementary Table S2. SARS-CoV-2 Sequence Variants Identified by NGS in Patient Material and Passage 2 of Virus Isolates.**

*^a^* SARS-CoV-2 identifier (ID; strain Switzerland/ZH-UZH-IMVxx/2020) and date of patient sample collection (dd/mm/2020).

*^b^* SARS-CoV-2 lineage as defined by Rambaut *et al.*, bioRxiv, 2020 and determined by CoV-GLUE (<http://cov-glue.cvr.gla.ac.uk>). SARS-CoV-2 clade as determined by Nextstrain (<https://clades.nextstrain.org/>).

*^c^* Difference from reference sequence (SARS-CoV-2/Wuhan-Hu-1; NC_045512.2) as determined by CoV-GLUE (<http://cov-glue.cvr.gla.ac.uk>). Bold amino-acid variants indicate that the variant was also detected in the original patient material that was directly sequenced.

*^d^* Additional variant in population at ≥ 15% frequency.

*^e^* Substitutions at these positions may change putative uncharacterized ORFs in overlapping coding regions (Gordon *et al.*, Nature, 2020; Firth, J Gen Virol, 2020).
