## Supplemental Table 3 for "SARS-CoV-2 Variants Reveal Features Critical for Replication in Primary Human Cells"

| **ID (Date)*^a^*** | **Lineage or Clade*^b^*** | **Amino acid variants*^c^*** | | | | | | | | **SNPs*^c^*** |
| --- | --- | --- | --- | --- | --- | --- | --- | --- | --- | --- |
|  |  | **Non-structural proteins** | **S** | **Orf3a** | **E** | **M** | **Orf6** | **Orf7a** | **N** |  |
| **BavPat1**  (2020) | B |  | S247R/S*^d^*  D614G  R682R/W*^d^* |  |  |  |  |  |  | C3037T, A11728G, T22303G, A23403G, C23606T |
| **IMV1**  (10/03) | B.2.2 / 19A | nsp2 (H237R)  nsp2 (D582A)  nsp6 (L37F) | D215N/D*^d^*  Δ677-688*^d^* | G251V |  |  | I14I/V*^d^* |  |  | A1515G, A2550C, C9223T, C11074T, G11083T, C14805T, T17247C, G22205A, G26144T, A27241G |
| **IMV2**  (09/03) | B.1 / 20C | nsp2 (T85I)  nsp12 (P323L)  nsp14 (P203P/L)*^d^* | D614G | Q57H*^e^* |  |  |  |  |  | C1059T, C3037T, C14408T, C18647T, A23403G, G25563T |
| **IMV3**  (11/03) | B.2 / 19A | nsp2 (P624P/L)*^d^*  nsp6 (L37F) | L5L/F*^d^*  F79L/F*^d^*  I233I/V*^d^*  H245H/R*^d^* | G251V |  | L17L/F*^d^* |  |  |  | C2676T, G11083T, C14805T, T17247C, T19146G, C21575T, T21797C, A22259G, A22296G, C25626T, G26144T, C26571T |
| **IMV4**  (11/03) | B.1 / 20A | nsp12 (P323L) | D614G |  | S6W |  |  |  |  | C3037T, C5167T, C14408T, C15324T, A23403G, C26261G |
| **IMV5**  (11/03) | B.1.1 / 20B | nsp12 (P323L) | D614G |  | S6S/L*^d^*  L19H/L*^d^* | T175M |  |  | R203K  G204R | C3037T, C14408T, T14646G, A23403G, C26261T, T26300A, C27046T, G28881A, G28882A, G28883C |
| **IMV6**  (29/03) | B.1.5 / 20A | nsp3 (K1693N)  nsp12 (P323L) | D614G |  | V5G |  |  |  | S194L*^e^* | C3037T, G7798T, C14408T, A20268G, A23403G, T26258G, C28854T |
| **IMV7**  (29/03) | B.1 / 20A | nsp12 (P323L) | G72R  D614G |  |  |  |  |  |  | C2638A, C3037T, C14408T, C15324T, C21691T, G21776A, A23403G |
| **IMV8**  (30/03) | B.1 / 20C | nsp2 (T85I)  nsp12 (P323L) | W64R/W*^d^*  S514F  D614G | Q57H*^e^* |  |  |  |  |  | C1059T, C3037T, C14408T, T21752C, C23103T, A23403G, G25563T, C29535T |
| **IMV9**  (07/04) | B.1 / 20C | nsp2 (T85I)  nsp12 (A97A/V)*^d^*  nsp12 (P323L)  nsp12 (S647I) | D614G | Q57H*^e^* |  |  |  |  |  | C1059T, C3037T, C13730T, C14408T, G15380T, A23403G, G25563T |
| **IMV10**  (14/04) | B.1.1 / 20B | nsp4 (S336L)  nsp12 (P323L) | S247R  D614G |  |  |  |  |  | R203K  G204R  I292T | C2462T, C3037T, C9561T, C14408T, T15837C, A22301C, A23403G, G28881A, G28882A, G28883C, T29148C |
| **IMV11**  (04/05) | B.1.5 / 20A | nsp3 (T970M)  nsp12 (P323L) | E583D  D614G |  | V5A |  |  |  | S194L*^e^* | C3037T, C5628T, A9529T, C14408T, A20268G, G23311T, A23403G, T26258C, C28854T |
| **IMV12**  (04/05) | B.1.1 / 20B | nsp12 (P323L) | H245H/R*^d^*  D614G  Δ675 |  | L37H/L*^d^* |  |  |  | R203K  G204R | C3037T, T11449C, C14408T, A20979G, A22296G, C22987T, A23403G, T26354A, G28881A, G28882A, G28883C |
| **IMV13**  (04/05) | B.1 / 20A | nsp6 (L37F)  nsp12 (P323L) | D614G |  | V5I/V*^d^* |  |  | H47N | R203K/R*^d^* | C3037T, G11083T, C14408T, C15324T, A23403G, G26257A, C27532A, G28881A |
| **IMV14**  (04/05) | B.1 / 20A | nsp2 (I273L)  nsp3 (K1083R)  nsp8 (A3V)  nsp12 (P323L) | D614G  Δ679-685*^d^* |  |  |  |  |  | L139F | C556T, A1622C, C3037T, A5967G, C12099T, C14408T, C15324T, A23403G, G28690T |

**Supplementary Table S3. SARS-CoV-2 Sequence Variants Identified by NGS in Working Stocks (Passage 3) of Virus Isolates.**

*^a^* SARS-CoV-2 identifier (ID; strain Switzerland/ZH-UZH-IMVxx/2020) and date of patient sample collection (dd/mm/2020).

*^b^* SARS-CoV-2 lineage as defined by Rambaut *et al.*, bioRxiv, 2020 and determined by CoV-GLUE (<http://cov-glue.cvr.gla.ac.uk>). SARS-CoV-2 clade as determined by Nextstrain (<https://clades.nextstrain.org/>).

*^c^* Difference from reference sequence (SARS-CoV-2/Wuhan-Hu-1; NC_045512.2) as determined by CoV-GLUE (<http://cov-glue.cvr.gla.ac.uk>).

*^d^* Additional variant in population at ≥ 15% frequency.

*^e^* Substitutions at these positions may change putative uncharacterized ORFs in overlapping coding regions (Gordon *et al.*, Nature, 2020; Firth, J Gen Virol, 2020).
